# Groucho running reveals disparate results between ground reaction force and tibia-fibula bone strain in runners

**DOI:** 10.64898/2026.08.14.744758

**Authors:** Arash Khassetarash, W. Brent Edwards

## Abstract

The relationship between external forces and bone strain in running is often complex and nonintuitive. We used Groucho running (i.e., running with exaggerated knee flexion) as a model to dramatically reduce the vertical ground reaction force (VGRF) and examined the relationship between peak VGRF and finite element (FE)-predicted tibia-fibula bone strain. Nine physically active males ran on an instrumented treadmill at 2.8 m/s with their preferred running technique, increased knee flexion (Groucho), and exaggerated knee flexion (Ex Groucho) in a randomized order. Strains at the tibia-fibula midshaft were calculated using computed-tomography-based FE modeling with loads and boundary conditions calculated from an inverse-dynamics based musculoskeletal model. Pressure-modified von Mises strain was used to quantify the peak strain (90^th^ percentile strain) and strained volume (volume of bone experiencing strains above 3000 µε). We further explored the relationship between peak VGRF, lower leg angle, and FE-predicted strain variables. The results showed that a 15.8% and 22.9% reduction in VGRF during Groucho and Ex Groucho, respectively, had no significant effect on FE-predicted peak strain (*p* > 0.304) and strained volume (*p*>0.053). Changes in peak VGRF did not correlate with FE-predicted strain variables (*p*>0.54) while changes in lower leg angle in the sagittal plane were moderately correlated (*r*>0.65; *p*<0.047). Our findings suggest that reductions in peak external forces do not always coincide with reductions in bone strain, especially in cases where running kinematics are dramatically altered. This work has important implications for designing gait retraining interventions based on reductions in external force measures.

## Introduction

Stress fractures are common overuse injuries in runners that frequently occur at the tibia ^1,2^. Mechanical fatigue is believed to play an important role in the development of these injuries ^3^, whereby cyclic loading results in bone failure at loading magnitudes well below the monotonic failure strength. Bone failure is ultimately strain controlled ^4^, and the relationship between bone strain and fatigue life (i.e., number of cycles to failure) is described by an inverse power law ^5,6^. The slope of the inverse power law is such that small changes in strain magnitude dramatically influence the fatigue life of bone. Accordingly, an accurate understanding of bone strain *in vivo* is of substantial importance for predicting the risk of stress fracture.

Quantifying bone strain *in vivo* is a non-trivial task. Direct measurement requires surgical attachment of strain gauges to the bony surface ^7,8^, whereas indirect estimation typically relies on numerical methods, such as combined musculoskeletal and finite element (FE) modeling ^9,10^. In contrast, quantifying external loads (i.e., ground reaction forces) is relatively straightforward owing to force platform data that are readily obtainable in a laboratory setting. Wearable technologies, including inertial measurement units and pressure-sensing insoles, have further expanded the ability to estimate external loads outside of the laboratory, primarily through statistical and machine-learning approaches to predict ground reaction force characteristics ^11,12^.

Because the assessment of bone strain remains difficult in practice, many rehabilitation and injury-prevention interventions have focused on reducing the magnitude of external loading as a surrogate measure of bone strain ^12–14;^ however, the relationship between bone strain and external load is often complex and nonintuitive. Indeed, our previous work illustrated only weak, insignificant correlations (i.e., *R*^2^=0.233; p=0.285) between changes in peak vertical ground reaction force (VGRF) and FE-predicted bone strain when manipulating flight time during running ^15^. From a multibody dynamics perspective, the orientation of the lower-extremity segments can substantially influence how external loads are balanced by internal forces and moments. (e.g., ^16^; The geometry and material properties of bone will also dramatically influence bone strain resulting from these internal loads.

We postulate that certain kinematic patterns within a runners’ capability can modify internal loading and bone strain independent of changes in external force magnitude. Groucho running, first described by McMahon et al. (1987), is a running technique characterized by exaggerated knee flexion and reduced leg stiffness, and substantial reduction in peak VGRF. As expected, this exaggerated knee flexion requires more muscle activity that dramatically increases the metabolic cost of running (up to 50%; ^17^. Higher muscle forces elevate internal loading, which would be expected to increase bone strain. Thus, the purpose of this study was to examine the relationship between peak ground reaction force and FE-predicted bone strain at the tibia-fibula bone during Groucho running. We hypothesized that lowering the peak VGRF in Groucho running would be associated with higher bone strain when compared to preferred running. We further hypothesized that tibia-fibula strains would be better predicted by lower leg angle at midstance than VGRF.

## Methods

### Groucho running: mechanics and definitions

Groucho running created an opportunity to manipulate peak VGRF by modulating the knee flexion angle during running. We postulate that the minimum theoretical peak VGRF (*F_max_*) by Groucho running can be calculated from the impulse-momentum relationship and solving for a theoretical zero vertical contact velocity (see Appendix A for proof):

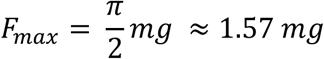

where *m* is the body mass and *g* is the gravitational constant. In practice, having zero vertical velocity at contact is difficult and unsustainable. Instead, we used this calculation as a guide to determine a percentage reduction in peak VGRF for our intervention. An average peak VGRF for a person running at 10 km/h is ∼2.5 *mg* ^18^. Therefore, lowering the peak VGRF by 25% would push the peak VGRF to 1.87 *mg*, which is still comfortably higher than the theoretical limit. We attempted to reduce peak VGRF using two interventions; Groucho running: hereafter called “Groucho” and exaggerated Groucho: hereafter called “Ex Groucho”. For Groucho and Ex Groucho, the runners attempted to create a peak VGRF that was 85% and 75% of their peak VGRF during preferred running, respectively.

### Participants

Previous gait modification studies of a similar nature (e.g., stride manipulation) have reported large effect sizes (*Co*ℎ*en*’*s d* = 1.25 − 2.75) for internal and external loading variables (Bonnaerens et al., 2021, 2019). An *a priori* power analysis was performed in G*Power (v3.1.9.7) using a conservative effect size corresponding to *Co*ℎ*en*’*s d* = 1.2 (equivalent to Cohen’s *f* = 0.6 for repeated-measures ANOVA), *α* = 0.05, statistical power of 1 − *β* = 0.80, three repeated measures, and an assumed correlation of *r* = 0.50 among repeated measures. The analysis indicated that a minimum sample size of N = 8 participants was required. We aimed to recruit a conservative convenience sample of 10 physically active participants, but recruitment was hindered by the COVID-19 pandemic, yielding N = 9 males (age: 27.2 ± 1.83 years, height: 1.76 ± 0.09 m, body mass: 75.5 ± 11.5 kg); their minimum weekly running milage was 10 km. The participants had no injuries within a six-month period prior to the data collection, and the study protocol was approved by our local Ethics committee (REB #19–1845) and all the participants provided written informed consent.

The participants visited the lab on 3 occasions. On the first visit, participants were familiarized with running on a split-belt instrumented treadmill (Bertec Corp., Columbus, OH) and the Groucho running technique. We asked the participants to run at a target speed of 2.8 m/s using their preferred technique while we collected VGRF for 5 minutes. Groucho running is expected to increase the metabolic cost of running by up to 50% ^17^, therefore, the speed was intentionally chosen conservatively to assure feasibility for an average physically active individual. We then calculated the average peak VGRF during the last 30 seconds and called it *Fmax*. After a sufficient rest period, a live view of the VGRF was projected on a screen in front of the participant while they ran on the instrumented treadmill. A target ribbon was overlayed on top of the VGRF graph. The participant was then instructed to keep their peak VGRF within the target ribbon by “exaggerating their knee flexion during stance”. The target ribbon was set to 85 ± 2.5% and 75 ± 2.5% of *Fmax* for Groucho and Ex Groucho, respectively. All participants were able to successfully modify their technique within the first 2 minutes of running.

During the second visit, lower-extremity and pelvic kinematics were tracked using 20 retroreflective markers placed on the left leg and pelvis. Marker clusters consisted of three markers on the foot, four on the shank, four on the thigh, and three on the pelvis. Additional anatomical markers were placed on the medial and lateral malleoli, the medial and lateral femoral epicondyles, and the left and right greater trochanters to define the ankle, knee, and hip joint centers, respectively. A static trial was recorded to establish segmental reference frames. The participant then warmed up by running on the treadmill for 5 minutes at their preferred speed. The participant then ran at 2.8 m/s using their preferred technique (Preferred), Groucho, and Ex Groucho for 5 minutes in a randomized order. Sufficient rest was provided between trials *ad libitum* to avoid exhaustion. At the 2.5-minute mark of each trial, thirty seconds of three-dimensional external forces and moments were recorded at 2000 Hz and motion capture data were recorded at 200 Hz (Vicon Motion Analysis Ltd., Oxford, UK). Like the familiarization session, the participant targeted 85% and 75% of *Fmax* guided by a ribbon on the live VGRF traces during Groucho and Ex Groucho, respectively.

The third visit was dedicated to computed tomography (CT) imaging. A CT scan capturing the entire tibiofibular complex was obtained for each participant using a GE Revolution GSI (GE Healthcare, Chicago, IL). A scan protocol of 120 kVp and 180 mA was used, and images were reconstructed with a 0.486 mm × 0.486 mm in-plane pixel-size and a 0.625 mm slice thickness. A calibration phantom with known hydroxyapatite densities (QRM GmbH, Mohrendorf, Germany) was used to convert CT attenuation values to bone mineral equivalent density (ρHU).

### Musculoskeletal modeling

A previously established inverse dynamics-based musculoskeletal modeling pipeline ^9,21^ was used to calculate internal loads. First, a system of rigid bodies was created for the foot, lower leg, thigh, and pelvis segments using the marker data. Then, a Cardan sequence of flexion-extension, abduction-adduction, and internal-external rotation was used to extract joint angles with respect to the neutral configuration. Kinetic data from the instrumented treadmill were first down-sampled to 200 Hz and then used as inputs to solve for net joint reaction forces and moments. A musculoskeletal model comprised of 44 muscle-tendon units ^22^ was used for simulations performed for every 1% of the stride with an objective function to minimize the sum of muscle stresses squared ^23^:

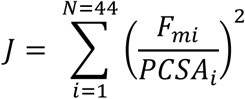

where *F_mi_* and *PCSA_i_* are the force and physiological cross-sectional area of the *i*th muscle, respectively. The net joint moments and maximum muscle forces (adjusted by force-velocity and force-length) were used as equality and inequality constraints, respectively. The joint moment constraints included the ankle and knee flexion-extension moments as well as the hip abduction-adduction and flexion-extension moments.

### Finite element modeling

CT images of the left tibia and fibula were manually segmented using the Mimics Innovation Suit (v21 Materialize, Leuven, Belgum) and converted to a FE mesh of the tibia-fibula in NetGen/NGSolve (https://ngsolve.org ). An average of 300,000 and 50,000 quadratic tetrahedral elements were used to model the tibia and fibula bones, respectively. A convergence analysis was previously performed with this FE model, which demonstrated that increasing element number from 130,000 to 270,000 changed peak strains by 4% ^24^. Bone was modelled as an inhomogeneous orthotropic material with elastic moduli being a function of apparent density (*ρ_app_*):

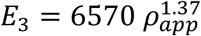

where *E*_3_ is the elastic modulus in the longitudinal direction and *ρ_app_* is equal to 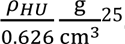. The elastic and shear moduli in the medio-lateral and anterior-posterior directions were defined as a function of *E*_3_ according to Rho et al., (1995):

*E*_1_ = 0.574 *E*_3_; *E*_2_ = 0.577 *E*_3_; *G*_12_ = 0.195 *E*_3_; *G*_23_ = 0.265 *E*_3_; *G*_31_ = 0.216 *E*_3_; *υ*_12_ = 0.427; *υ*_23_ = 0.234; *υ*_31_ = 0.405

where *G* is the shear modulus and *υ* is the Poisson’s ratio with subscripts 1, 2, and 3 denoting the medio-lateral, anterior-posterior, and longitudinal directions, respectively. These material constants have been shown to produce excellent agreement in FE-predicted strains of cadaveric human tibiae ^27^.

### Boundary conditions

The tibial plateau was fixed in all degrees of freedom with an additional element on the medial proximal tibia fixed in the anterior-posterior direction. The ankle joint was fixed in the anterior-posterior and medio-lateral directions. Adjacent elements of the tibia and fibula at the proximal and distal ends were bound together. Additionally, anterior and posterior proximal tibiofibular ligaments were modelled by spring elements with stiffness according to Marchetti et al. ^28^. Ankle joint contact force and a residual moment ^10^ were applied to the ankle joint. Forces from sixteen lower extremity muscles (Semimembranosus, Semitendinosus, long and short head of Biceps Femoris, Sartorius, Gracilis, Soleus, Tensor Fascia Latae, Tibialis Anterior, Tibialis Posterior, Flexor Digitorum Longus, Flexor Hallucis Longus, Peroneus Longus, Peroneus Brevis, Peroneus Tertius, and Extensor Digitorum Longus), along with patellar tendon force were modelled as point forces at their insertion on the tibia and fibula. This FE analysis pipeline was previously verified ^29^ using experimental data from *in vivo* bone pin ^30^ and strain gauge measurements ^7^. The FE simulations were carried out in Abaqus (version 2021, Dassault System, RI, USA) using muscle and joint contact forces at the moment of peak ankle joint contact force. We calculated pressure-modified von Mises strains within the diaphysis of the tibia and fibula (mid 60% of the tibial length) and extracted the 90^th^ percentile strain as a measure of peak strain. We also extracted a measure we call “strained volume” (i.e., the volume of the bone experiencing strains above a 3000 µε threshold), as this measure was previously shown to correlate strongly with the fatigue life of whole bone ^6^.

### Statistical Analysis

All statistical analyses were conducted in R Studio (2023.12.1) with the criterion alpha-level set to 0.05. To test the primary hypothesis that Groucho running increases tibia-fibula strains, separate repeated measures ANOVAs were performed for each strain variable. Running condition (three levels) was treated as a within-subject factor. When a significant main effect of condition was observed, pairwise comparisons between conditions were conducted using paired t-tests with Holm-adjusted *p*-values to control for family-wise error. Because two primary dependent variables were analyzed, *p*-values for the main effects were additionally adjusted using the Holm procedure.

Spatiotemporal variables, peak VGRF, peak resultant ankle and knee joint contact forces were analyzed using separate repeated-measures ANOVAs to verify that the running techniques produced distinct movement patterns. These analyses were considered secondary and were interpreted as supportive of the primary analysis. To account for multiple comparisons among the secondary outcome variables, p-values from the omnibus repeated-measures ANOVAs were adjusted using the Benjamini–Hochberg false discovery rate procedure ^31^.

To address our second hypothesis, change scores (Δ) were calculated for each participant relative to the reference condition (Preferred). Data from the two Groucho conditions (Groucho and Ex Groucho) were pooled, and repeated-measures correlation was used to assess the within-subject association between ΔVGRF, Δleg angle, and Δstrain variables (peak strain and strained volume), accounting for non-independence of observations within participants.

For all the variables and tests, normality of residuals was assessed using the Shapiro–Wilk test. Sphericity was evaluated using Mauchly’s test; when violated, Greenhouse–Geisser corrections were applied. Effect sizes were also reported as the partial eta squared (η^2^) from the ANOVA.

## Results

Time histories of the VGRF during the three running techniques are presented in **Figure 1** for a representative runner. Attempts to decrease the peak VGRF using Groucho running was successful. Overall, a significant effect of running condition on the peak VGRF was observed (*F*(2, 16) = 123.79, *p* <0.001, η^2^ =0.361; **Figure 2A**). Mean (± SD) peak VGRF values were 1757 ± 214 N (Preferred), 1479 ± 210 N (Groucho), and 1354 ±190 N (Ex Groucho). Pairwise comparisons indicated that VGRF was significantly lower in both Groucho and Ex Groucho (by 15.8 % and 22.9 %, respectively) compared to Preferred (both *p* < .001) and further reduced (by 8.5 %) in Ex Groucho compared to Groucho (*p* < .001).

**Figure 1.**
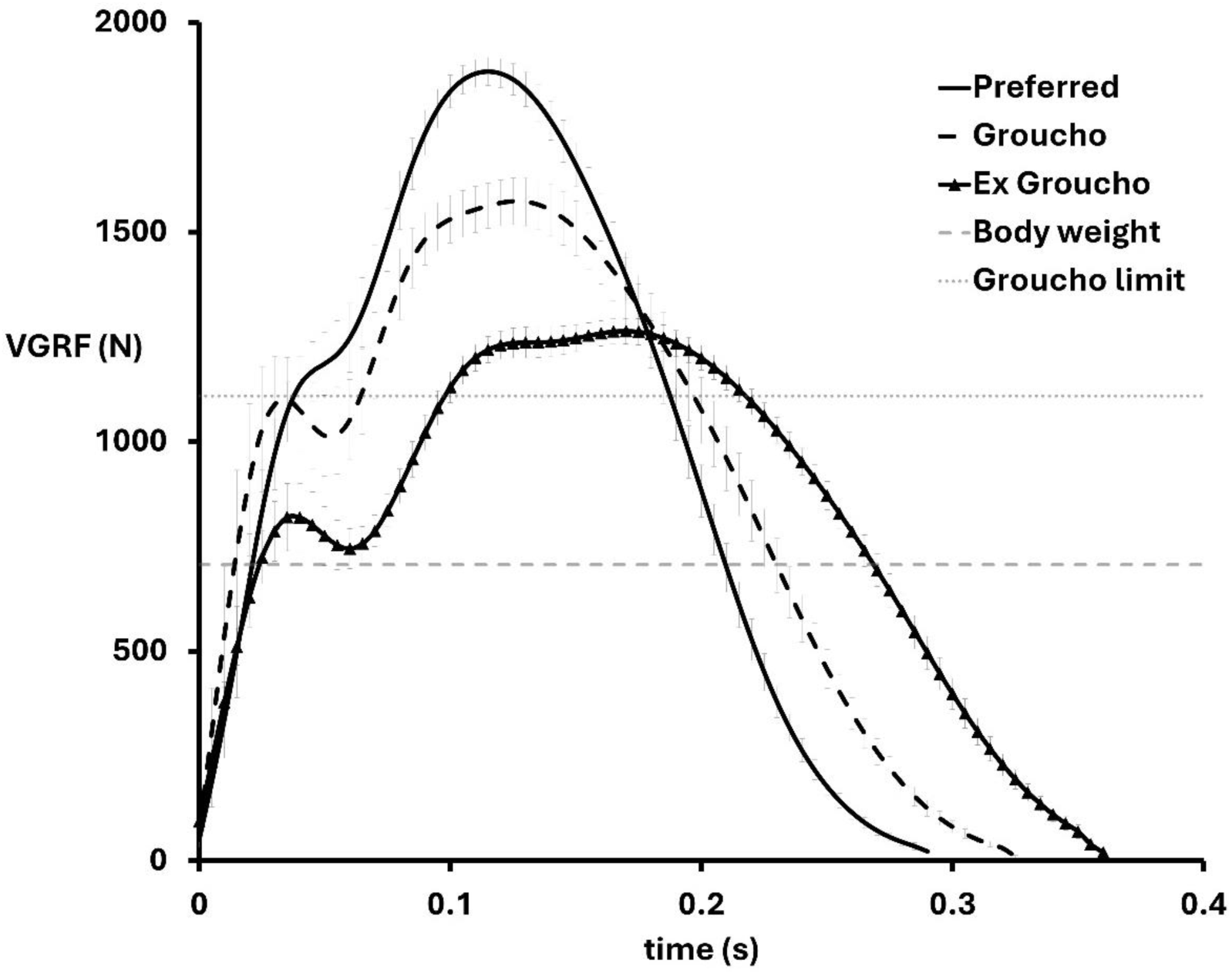
Time history of average and standard deviation (of 10 stances) vertical ground reaction force (VGRF) for a representative participant during preferred (black solid line), Groucho (black dashed line) and Ex Groucho (black solid line with triangle markers). The participant’s body weight (dashed gray line) and theoretical lower limit (see appendix A) of peak VGRF during Groucho running (dotted gray line) are also shown.

**Figure 2.**
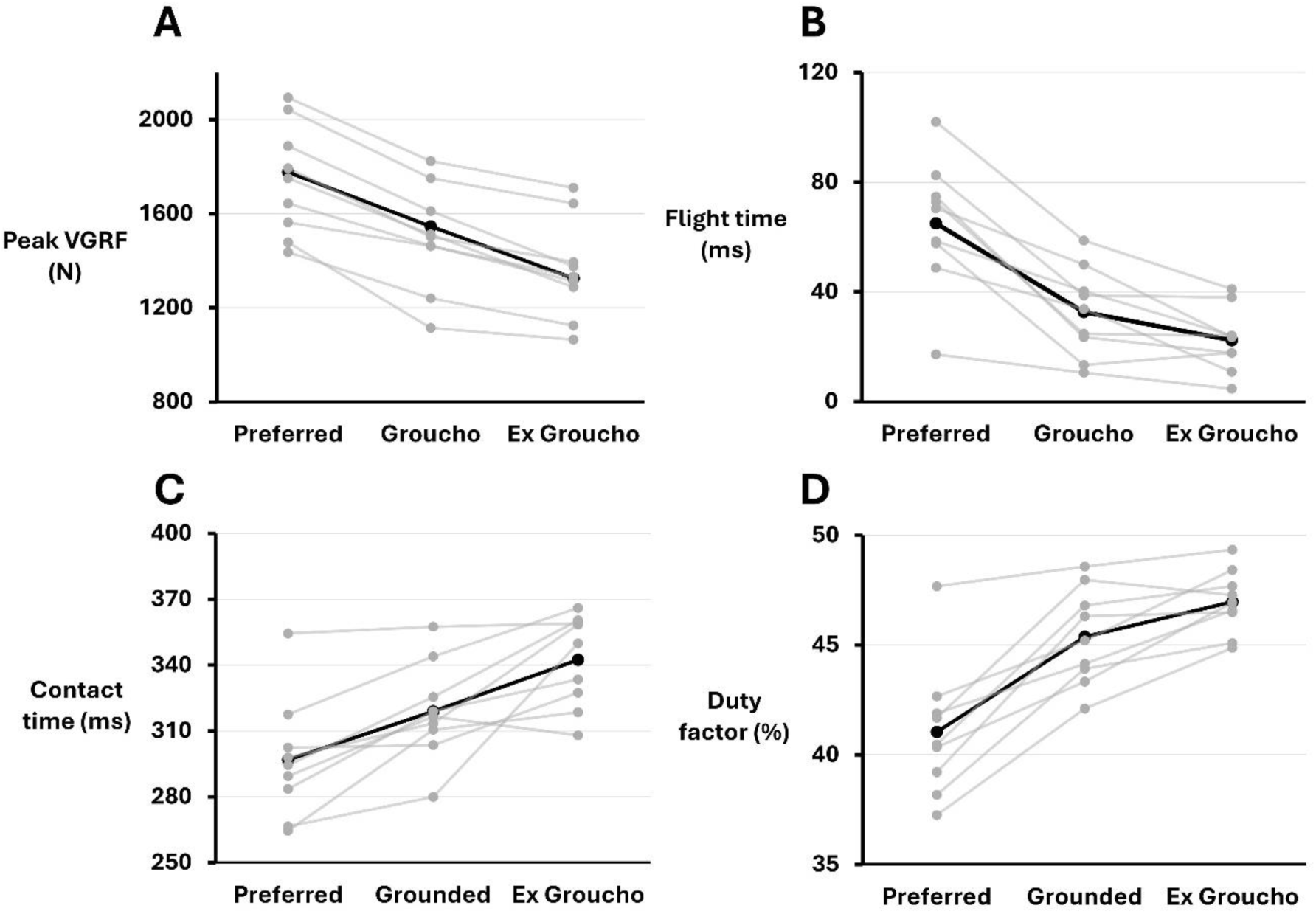
Peak Vertical ground reaction force (VGRF) and spatiotemporal variables during the three running conditions: A) peak VGRF B) Flight time, C) contact time, and D) duty factor. In all graphs, the thick black line represents the average of all participants, and the gray transparent lines represent individual participants.

A significant effect of running condition on flight time was observed (*F*(2, 16) = 44.00, *p* <0.001, η^2^ =0.537; **Figure 2B**). Mean (± SD) flight times were 64 ± 23 ms (Preferred), 33 ± 16 ms (Groucho), and 22 ± 12 ms (Ex Groucho). Pairwise comparisons indicated that flight time was significantly reduced (by 7.5 % and 15.4%, respectively) in both Groucho and Ex Groucho compared to Preferred (both *p* < 0.002) and further reduced (by 7.4%) in Ex Groucho compared to Groucho (*p* =0.019).

A significant effect of running condition on contact time was observed (*F*(2, 16) = 19.70, *p* <0.001, η^2^ =0.409; **Figure 2C**). Mean (± SD) contact time were 297 ± 27 ms (Preferred), 319 ± 22 ms (Groucho), and 342 ± 21 ms (Ex Groucho). Pairwise comparisons indicated that contact time was significantly higher (by 7.5 % and 15.4% respectively) in both Groucho and Ex Groucho compared to Preferred (both *p* < 0.002) and further increased (by 7.4%) in Ex Groucho compared to Groucho (*p* =0.019).

A significant effect of running condition on duty factor was observed (*F*(2, 16) = 48.55, *p* <0.001, η^2^ =0.567; **Figure 2D**). Mean (± SD) duty factor were 41 ± 3 % (Preferred), 45 ± 2 % (Groucho), and 47 ± 1 % (Ex Groucho). Pairwise comparisons indicated that duty factor was significantly higher (by 10.5 % and 14% respectively) in both Groucho and Ex Groucho compared to Preferred (both *p* < 0.001) and further increased (by 3.5%) in Ex Groucho compared to Groucho (*p* =0.012).

No significant effect of running condition on step frequency was observed (*F*(2, 16) = 2.02, *p* =0.165, η^2^ =0.071). Mean (± SD) step frequency were 166 ± 10.5 steps/min (Preferred), 171 ± 10 steps/min (Groucho), and 165 ± 12 steps/min (Ex Groucho).

The distribution of pressure-modified von Mises strain (**Figure 3**) and strained volume (**Figure 4**) are shown for two representative participants. One participant (A) showed an increase in FE-predicted peak strains, and the other participant (B) showed a decrease in peak strains during both Groucho and Ex Groucho compared to Preferred. No significant effect of running condition on peak strains was observed (*F*(1.21, 9.68) = 1.24, *p* = 0.304, η^2^ =0.026). Mean (± SD) peak strain values were 4695 ± 861 µε (Preferred), 4339 ± 1014 µε (Groucho), and 4428 ±1078 µε (Ex Groucho). **Figure 5A**. In contrast, a significant effect of running condition on strained volume was observed (*F*(2, 16) = 5.127, *p* = 0.019, η^2^= 0.097). However, Holm-adjusted pairwise comparisons did not reveal significant differences between any conditions (all *p* ≥ 0.053). Mean (± SD) strained volume values were 10546 ± 3227 mm^3^ (Preferred), 8357 ± 2771 mm^3^ (Groucho), and 8956 ±2925 mm^3^ (Ex Groucho). **Figure 5B**.

**Figure 3.**
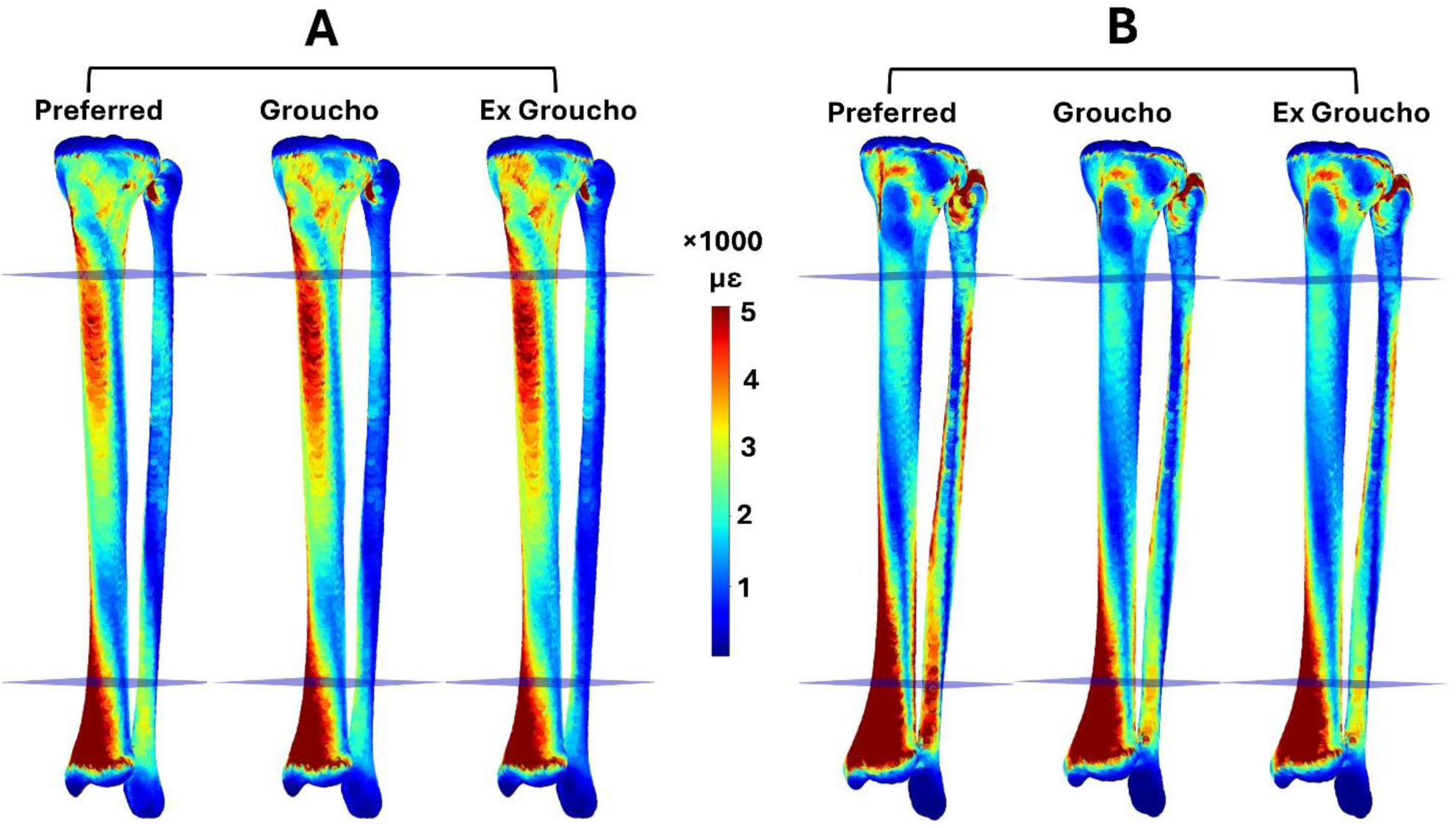
Distribution of pressure-modified von Mises strain for two representative runners (participants A and B). Runner A showed an increase in peak strain (4713, 4985, 4952 µε during Preferred, Groucho, and Ex Groucho, respectively) while runner B showed a decrease in peak strains (4744, 4081, 4239 µε during Preferred, Groucho, and Ex Groucho, respectively).

**Figure 4.**
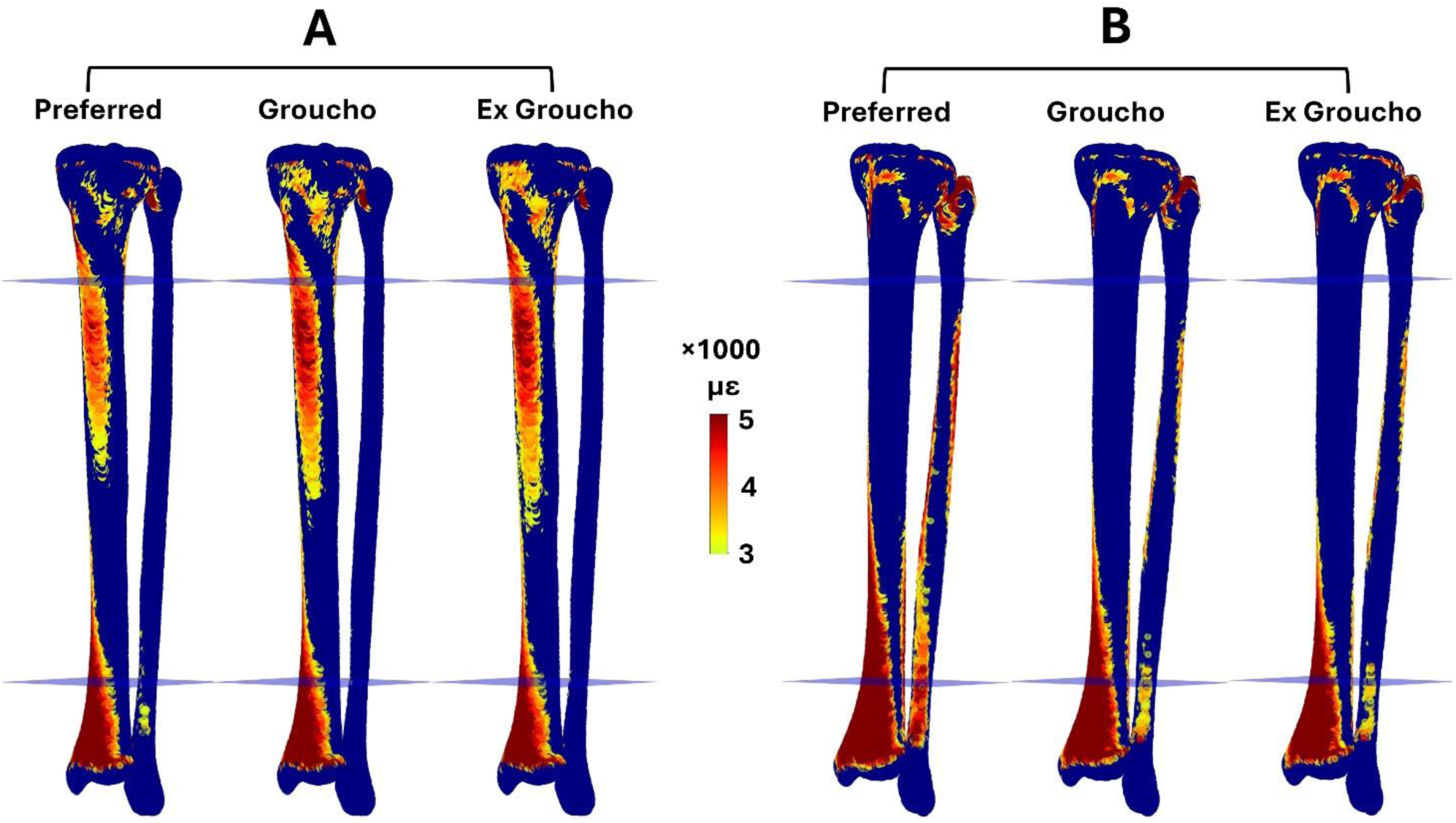
Strained volume (i.e., volume of bone experiencing strains above 3000µε) for two representative runners. Runner A showed an increase in strained volume during Groucho and Ex Groucho compared to preferred (9349, 10046, 10407 mm^3^ during Preferred, Groucho, and Ex Groucho, respectively), while runner B showed a decrease in strained volume (11135, 8672, 8871 mm^3^ during Preferred, Groucho, and Ex Groucho, respectively).

**Figure 5.**
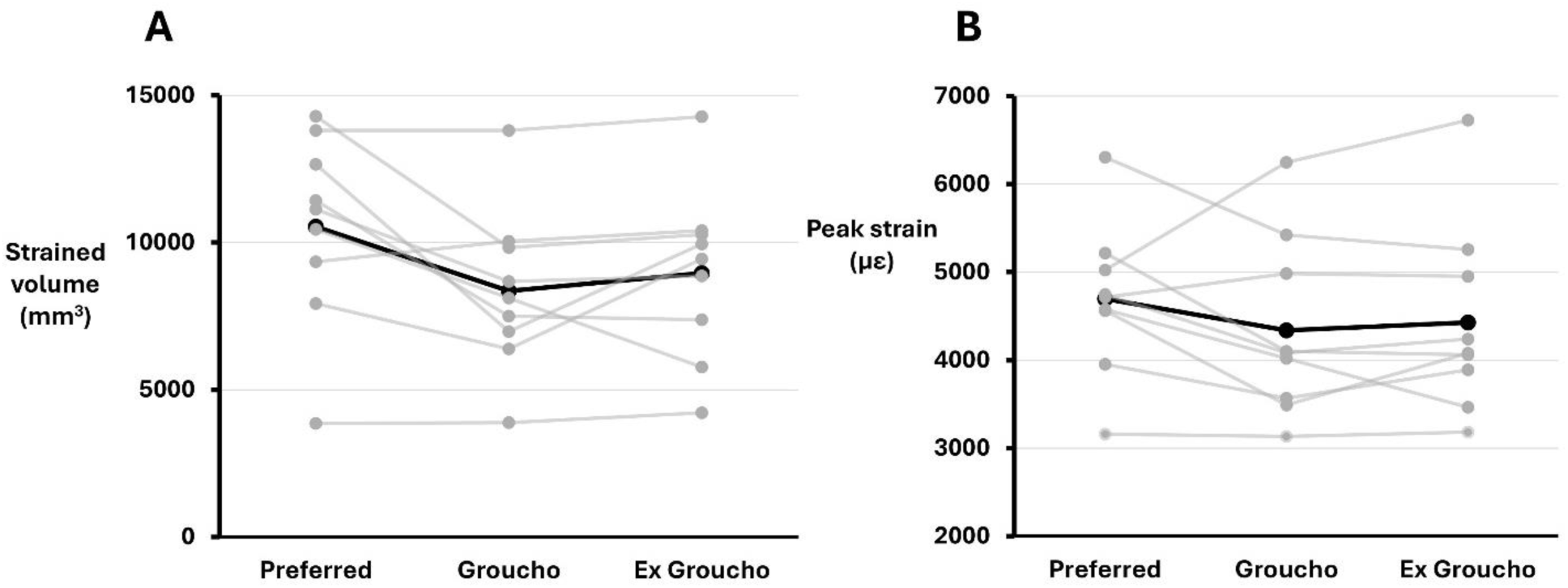
Finite element-predicted A) peak strain (i.e., 90th percentile strain within the midshaft of the tibia-fibula) and B) strained volume (i.e., volume of bone experiencing strains above 3000 με within the midshaft of tibia-fibula). In both graphs, the thick black line represents the average of all participants, and the gray transparent lines represent individual participants.

A significant effect of running condition was observed for the peak resultant ankle contact force (*F*(2, 16) = 40.99, *p* <0.001, η^2^ =0.277; **Figure 6A**). Mean (± SD) peak resultant ankle contact forces were 7327 ± 933 N (Preferred), 6400 ± 879 N (Groucho), and 6121 ± 836 N (Ex Groucho). Pairwise comparisons indicated that the peak resultant ankle contact force was significantly lower (by 12.6 % and 16.5 % respectively) in both Groucho and Ex Groucho compared to Preferred (both *p* < 0.001) and further decreased (by 4.4 %) in Ex Groucho compared to Groucho (*p* =0.037). In contrast, no significant effect of running condition was observed for the peak resultant knee contact force (*F*(1.1, 8.8) = 2.55, *p* =0.145, η^2^ =0.079; **Figure 6B**). Mean (± SD) peak resultant knee contact force were 9018 ± 1742 N (Preferred), 7646 ± 3025 N (Groucho), and 8941 ± 1849 N (Ex Groucho).

**Figure 6.**
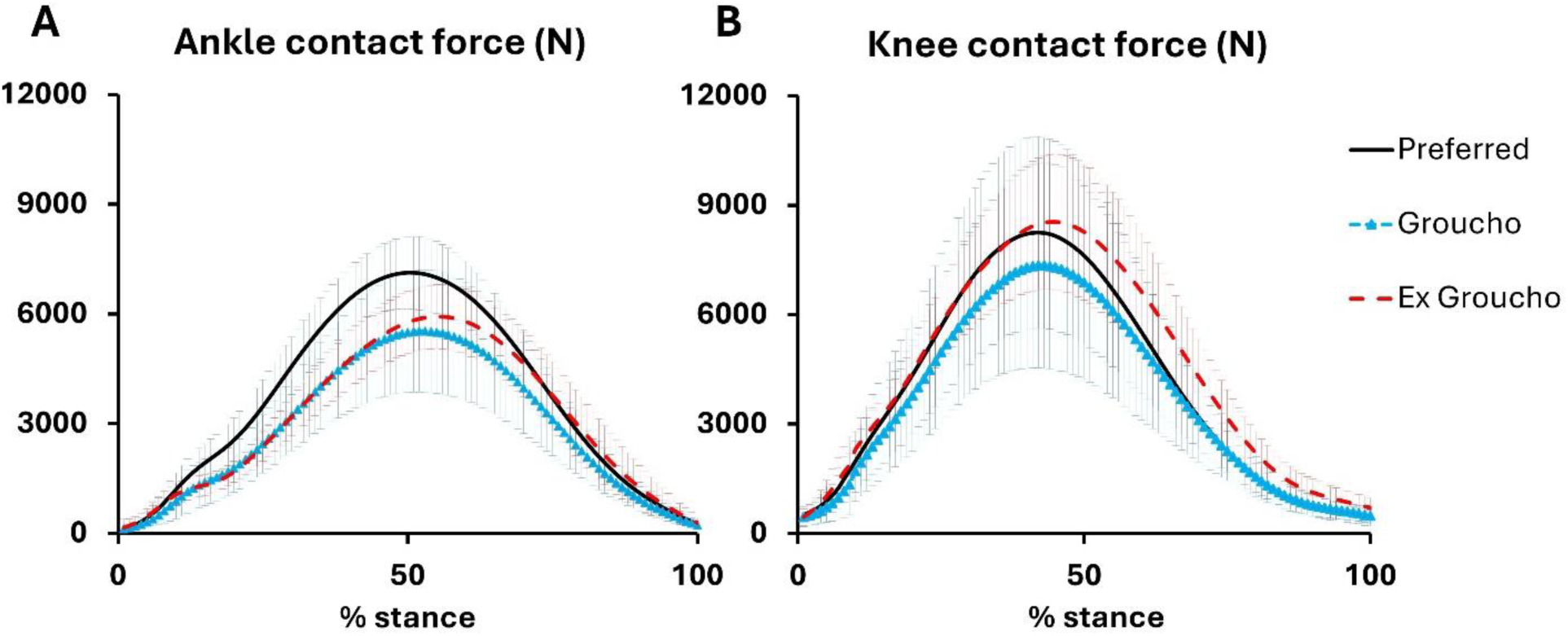
Ensembled average ±standard deviation (of 9 participants) of the resultant joint contact forces at the ankle (A) and knee (B) during the stance phase of running.

A repeated-measures correlation showed no significant association between change in peak VGRF and change in peak strain (*r*(8) = -0.216, p = 0.549, 95% CI [−0.744, 0.479]; **Figure 7A**). Similarly, no association was observed between the peak VGRF and strained volume (*r*(8) = -0.184, p = 0.611, 95% CI [−0.729, 0.504]; **Figure 7B**). In contrast, a significant positive association was found between the change in lower leg angle at midstance and change in peak strain (*r*(8) = 0.638, p = 0.047, 95% CI [0.014, 0.904]; **Figure 7C**). Similarly, a significant positive association was found between change in lower leg angle at midstance and change in strained volume (*r*(8) = 0.651, p = 0.041, 95% CI [0.037, 0.908]; **Figure 7D**).

**Figure 7.**
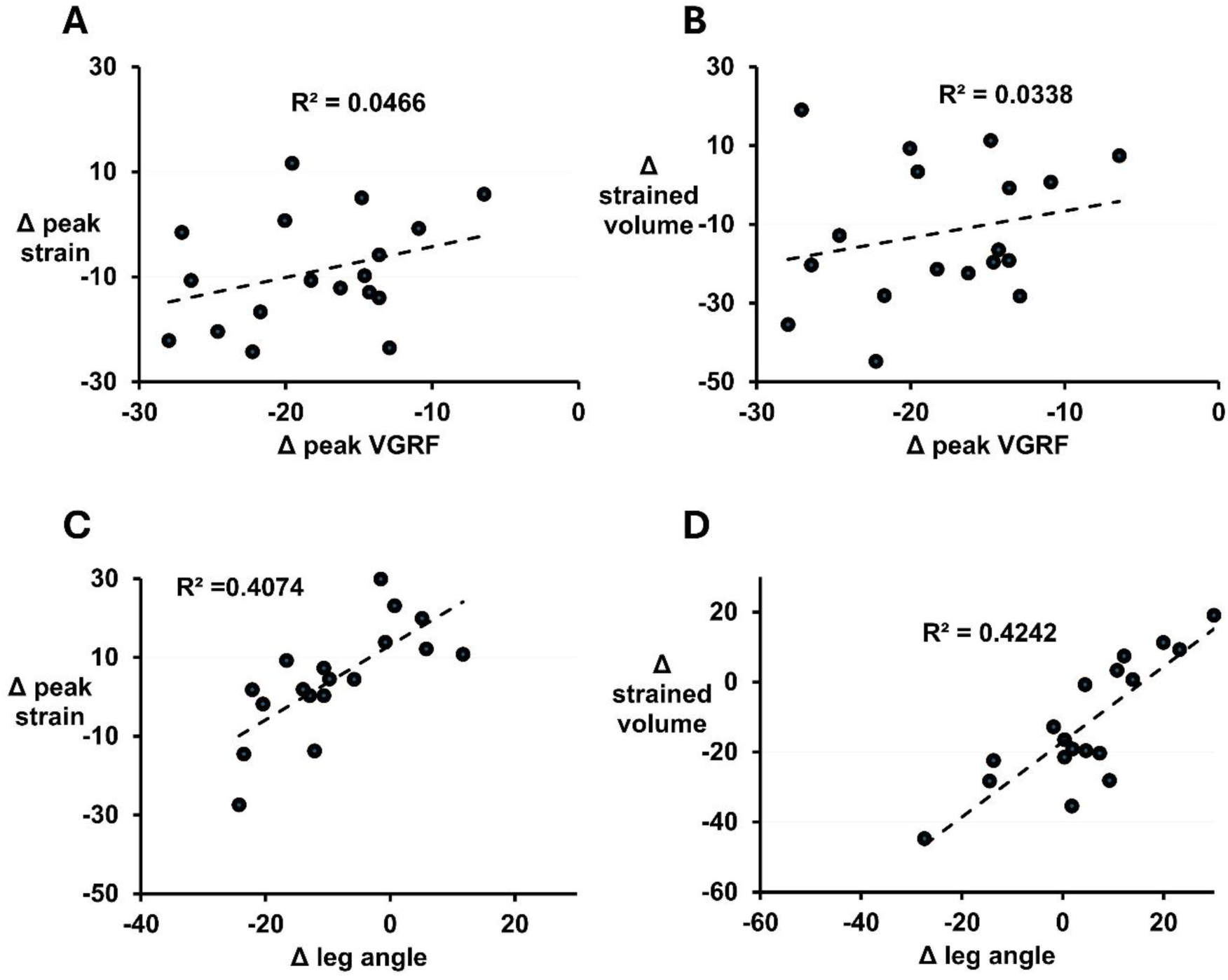
within-subject association between Δ peak strain and A) Δ leg angle, and B) Δ peak VGRF along with within-subject association between Δ strained volume and C) Δ leg angle, and D) Δ peak VGRF.

## Discussion

We used Groucho running (i.e., running with exaggerated knee flexion) as a tool to explore the complex relationship between the peak VGRF and tibia-fibula bone strain. We hypothesized that 1) decreasing peak VGRF by Groucho running would lead to an increase in tibia-fibula bone strains, and 2) changes in tibia-fibula bone strains would be better explained by changes in lower leg angle at midstance than changes in peak VGRF. Our results did not fully support our first hypothesis. In fact, neither the change in peak strains nor strained volume was statistically different during Groucho or Ex Groucho compared to preferred running, despite dramatic reductions in peak VGRF (≥15.8%). While some of the participants demonstrated increased strains during Groucho and Ex Groucho, others showed minimal change or reduced strains (**Figure 5**). On the other hand, our results supported the second hypothesis. There was no meaningful association between changes in peak VGRF and changes in FE-predicted strains (all *r* < 0.22, *p* > 0.54). In contrast, changes in the sagittal-plane lower leg angle at midstance explained approximately 40– 42% of the variance in FE-predicted strains (all *r* > 0.63, all *p* < 0.05).

There was large scatter in the FE-predicted peak strains (e,g., 3100 µε - 6300 µε during Preferred running) and strained volume for different participants. Similar variation between participants was previously reported by *in vivo* strain gauge studies ^7,8^, bone bending experimental measurements ^30^, and computational models ^10,24,32^. This large degree of scatter is expected given that the internal loading, geometry, and material properties of the bone can significantly affect bone deformation. In fact, the participant with the highest resultant knee contact force (∼11400 N) illustrated the highest peak strains (>6000 µε). These results clearly illustrate the complex and non-intuitive relationship among bone strain, internal loading, and external loading metrics.

Our findings can be explained using a simple force systems analysis, illustrating that an increase in lower leg angle during Groucho increases the moment arm of the ankle reaction force about the knee (Appendix B). This increased moment arm may offset the effect of lower ankle reaction force, resulting in marginal change — or potentially an increase— in the required knee extensor moment. In participants exhibiting greater knee extensor moments, this will require larger quadriceps and patellar ligament forces. Concomitantly, the tibia experiences a reduction in ankle contact force (and thus reduced plantarflexors, e.g., soleus force) due to reduced peak VGRF, which decreases the axial compressive component of loading. It is important to note that the tibia experiences bending as well as compressive loading during stance ^30^. A substantial reduction in axial loading due to decreased ankle joint contact force (e.g., decreased Achilles tendon force) can be offset by a relatively modest increase in knee contact force (e.g., increased patellar ligament force). Because the patellar ligament acts farther anterior to the centroid of the tibial diaphysis than the soleus attachment, increases in patellar ligament force are expected to preferentially increase sagittal-plane bending moments. Given that bending produces pronounced strain gradients and associated shear components across the cortex, increases in bending moments can lead to disproportionately larger increases in bone strain than comparable changes in axial loading. Therefore, even a dramatic decrease (by 22.9%) in peak VGRF may still be associated with minimal to no change in tibia and fibula bone strain.

Our previous work demonstrated that relying solely on external loading metrics may be insufficient for predicting bone strains and, consequently, the risk of stress fracture ^15^. The present findings are consistent with this observation, suggesting that, under certain conditions, changes in lower leg orientation may be more strongly associated with bone strain than external loading measures. It is important to note, however, that the observed association between sagittal-plane lower leg angle and FE-predicted strains should not be considered a general relationship applicable to all running conditions. Although Groucho running represents an extreme intervention, caution should be made when introducing gait retraining interventions targeting either a reduction in peak VGRF or increased duty factor^14,33^.

*In-vivo* measurement of bone strain during running is a non-trivial task given the complexity/uncertainty involved in direct strain gauge/bone pin measurements ^7,8,30^ and modeling approaches ^10,15,32^. This contrasts with the ease of measuring external loads using force plates and pressure insoles in the real world. Therefore, significant emphasis has been placed on measuring external loading such as ground reaction forces as a surrogate measure of bone strain. Especially with the widespread use of wearable technologies, many physics-based ^34^ and machine learning-based ^35,36^ methods have been developed to predict the VGRF during locomotion. More advanced methods in recent years focused on estimating peak longitudinal forces within the tibia ^37,38^ while ignoring the role of bending moments ^21,39,40^ on the development of tissue strains. The latter becomes increasingly important when the kinematics of running substantially deviate from a preferred average. Therefore, recommendations made by a hypothetical wearable device that measures and estimates risk of stress fractures would be erroneous in specific circumstances.

There are several limitations to this study. One potential concern is the uncertainty associated with forces predicted by musculoskeletal models. The neuromuscular strategy (i.e., cost function) for muscle force redundancy may have been changed as a result of changes in running technique. We used electromyography measurements from seven lower extremity muscles to examine the similarity between the model-predicted muscle forces and the EMG intensity (see Appendix C) across running conditions. We did not observe any differences in the similarities explored as a result of change in running technique (see Appendix C – Figures C1 and C2, and Tale C1). Therefore, the repeated-measures design likely minimized the influence of the limitation regarding the muscle force estimation, since each participant served as their own control across conditions. We also tested a relatively small sample of participants. Nevertheless, previous gait-manipulation studies ^15,19^ have demonstrated that similarly small sample sizes are sufficient to establish significant changes when meaningful relationships exist. Despite the limited sample size, we were still able to identify a significant relationship between lower-leg kinematics and FE-predicted strains. The ecological validity of Groucho running is also questionable. It is unlikely that humans would naturally adopt a Groucho running gait ^41^ because it is highly inefficient and metabolically demanding ^17^. However, the primary objective of this study was not to investigate the practicality of Groucho running itself, but rather to examine the relationship between peak VGRF, lower-leg kinematics, and FE-predicted tibia-fibula strains.

## Conclusion

In summary, our study demonstrated a non-intuitive relationship between the external loads and bone strain. The results showed that even a dramatic reduction in peak external loads of up to 22.9% can still cause an increase in internal loading if it is accomplished by significant changes in lower leg angle. This study highlights the importance of considering loading on the tissue of interest in gait retraining interventions rather than surrogate measures such as external loading.

## Acknowledgment

We acknowledge the support of the Natural Sciences and Engineering Research Council of Canada (RGPIN 02404-2021). We would like to thank Art Kuo, Jeremy Wong, Koen Lemire for their help in study design and implementation, and Mark Pineda for the help with data collection.

## Conflict of interest

No conflict of interest to declare.

## Notes

### Competing Interest Statement

The authors have declared no competing interest.

### Summary of Updates

Upon reviewing the generated PDF, noticed some of the hyperlinks to figures were not recognized properly. The new version has resolved this issue.

