## appendices for "Groucho running reveals disparate results between ground reaction force and tibia-fibula bone strain in runners"

### Appendix A

#### Proof of theoretical limit of peak vertical ground reaction force (VGRF) during running

Assume the initial vertical velocity of the centre of mass of a runner at ground contact is represented by $u$. By using impulse-momentum theory, the change in vertical velocity of the centre of mass during contact should be equal to the integral of the net force acting on the centre of mass. Therefore:

$$m\,\Delta u = \int_0^{tc} (F - mg)\ dt \tag{A1}$$

Where $m$ is the mass of the runner, $F$ is the VGRF, $g$ is the gravitational constant, $tc$ is the contact time, and $t$ denotes time. In a steady-state run, the vertical velocity of the centre of mass ($u$) will flip sign. Therefore:

$$2mu = \int_0^{tc} F dt - \int_0^{tc} mg\ dt \tag{A2}$$

$$u = \frac{1}{2m}\int_0^{tc} F dt - g\frac{tc}{2} \tag{A3}$$

We can model the VGRF during contact as a simple sine wave with a maximum and the period equal to twice the contact time:

$$F = F_{max}\sin\left(\frac{\pi t}{tc}\right) \tag{A4}$$

Therefore:

$$u = \frac{1}{2m}\int_0^{tc}\left(F_{max}\, si\, n\left(\frac{\pi t}{tc}\right)\right) dt - g\,\frac{tc}{2} \quad \text{(A5)}$$

And solving for the integral:

$$u = \frac{1}{2m}\left[-\frac{tc}{\pi}F_{max}\,\cos\left(\frac{\pi t}{tc}\right)\right]\Big|_0^{tc} - g\,\frac{tc}{2} \quad \text{(A6)}$$

For near-zero initial contact velocity ($u = 0$):

$$\frac{1}{2m}\left[-\frac{tc}{\pi}F_{max}\,\cos\left(\frac{\pi t}{tc}\right)\right]\Big|_0^{tc} = g\,\frac{tc}{2} \quad \text{(A7)}$$

$$\frac{2\,tc}{\pi}F_{max} = (mg)\,tc \quad \text{(A8)}$$

$$F_{max} = \frac{\pi}{2}(mg) \approx 1.57\,mg \quad \text{(A9)}$$

### Appendix B

### A simplified inverse dynamics analysis

For simplicity, we focused our attention on a two-dimensional analysis in the sagittal plane. We considered the forces and segmental configuration at midstance during running where the anterior-posterior component of the external force was zero and the external force was perfectly vertical. Error! Reference source not found. shows an inverse dynamics analysis for the ankle and the knee joint. (Note 1: for the purpose of this demonstration, we omitted the inertial terms as: 1) the influence of these terms are small and negligible compared to the external force contributions and 2) the inertial terms are not very different between the two running techniques). (Note 2: the inertial terms were fully considered in the 3D inverse-dynamics-based musculoskeletal modeling approach explained in the manuscript).

A significant reduction in the peak VGRF (shown as $F$) would cause a corresponding significant reduction in the ankle reaction force and reaction moment when the moment arm ($d$) remains constant (Error! Reference source not found.**A**). The reduction in ankle reaction moment equates to a reduction in plantarflexor forces such as soleus and consequently, a reduction in ankle joint contact force. While knee reaction force is also directly affected by the magnitude of the external force, the reaction moment is dependent on the angle of the lower leg segment with respect to the vertical axis (Error! Reference source not found.**B**). The larger this angle (and potentially the knee flexion angle), the larger the knee reaction moment. An increase in the knee reaction moment would likely amplify quadriceps, patellar tendon, and knee joint contact forces. Because the patellar tendon inserts anteriorly on the tibial tuberosity, if may alter tibial bending mechanics and contribute to increased tibial strain. From a mechanics of materials perspective, bending-induced strains on the bone surface are typically much greater than strains produced by purely axial (longitudinal) loading. Consequently, a reduction in axial compressive loading resulting from decreased soleus force can be offset, or even exceeded, by increased bending moments associated with greater patellar tendon force. Therefore, even a substantial reduction in VGRF may still lead to increased internal tibial loading if it is accompanied by a large increase in knee flexion angle.

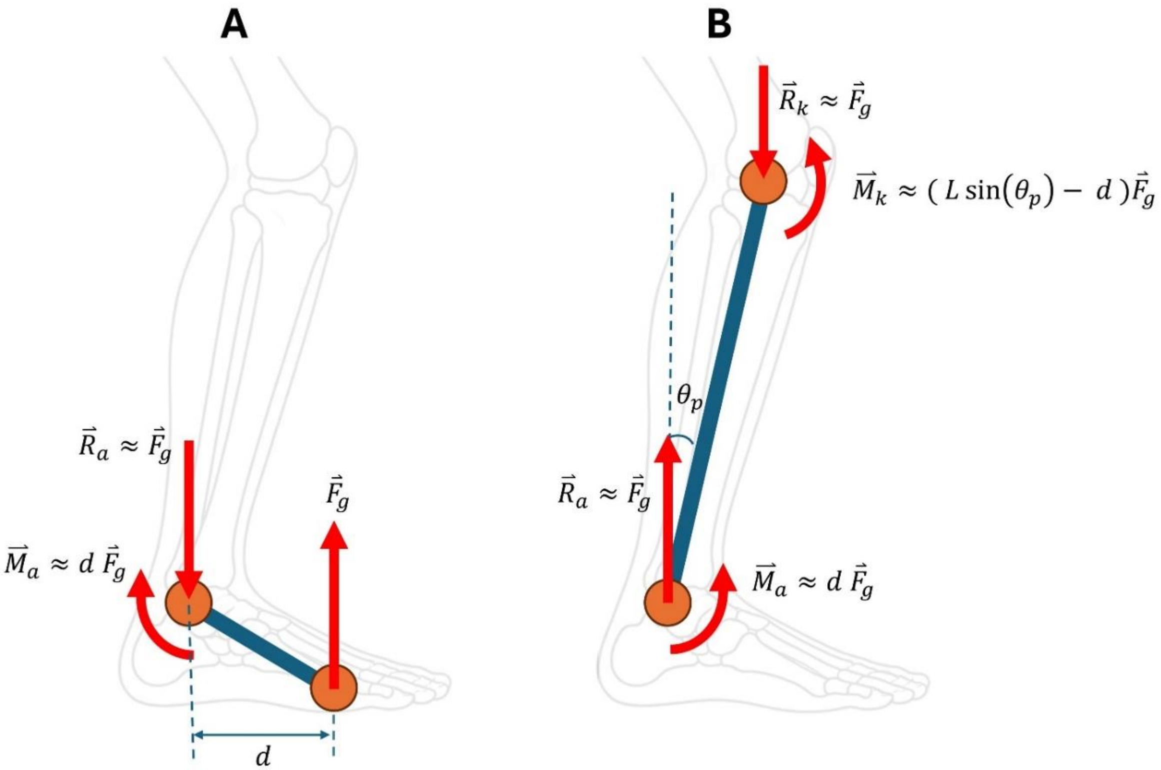


**Figure B1.** A schematic representation of 2D inverse dynamics analysis of the A) ankle reaction forces/moments and B) knee reaction forces/moments. Note that for simplicity, we have omitted the inertial terms.

### Appendix C

### Evaluating the agreement between muscle activity measured by electromyography and musculoskeletal-modeling predicted muscle forces

#### Introduction

Predicting muscle forces during locomotion is challenging given mathematical redundancy [1]. An infinite combination of muscle forces can be predicted to produce a required net moment about a joint. This problem is usually solved through optimization assuming a cost function [2]. In the current study, we used a cost function to minimize the square of muscle stresses [3]. While this cost function has been generally shown to predict agreeable muscle forces when compared to electromyography [4] and across a range of running speeds and grades [5], we wanted to assure that manipulations in running technique was not associated with significant alterations in agreement.

#### Methods

To evaluate the agreement between the forces predicted from musculoskeletal modeling and muscle activation, we collected EMG data from 7 muscles: Gastrocnemius Medialis, Soleus, Tibialis Anterior, Vastus Lateralis, Rectus Femoris, Biceps Femoris, and Gluteus Maximus. The skin was prepared by shaving, abrading, and using alcohol swipes prior to electrode attachments. Double-differential surface sensors (DE-2.1. Delsys Inc.) with electrode distance of 10 mm were placed over the muscle bellies based on SENIAM recommendations [6]. A reference electrode was placed on the patella. Signal quality was assured by visual inspection while the participant performed multiple contractions using the target muscles. EMG data was recorded at 2000 Hz and synchronized with kinetic and kinematic data.

EMG data were filtered using a 4th order, zero-lag, bandpass Butterworth filter with cut-off frequencies of 20 Hz and 500 Hz. The smoothed (using a 4th order, zero-lag, low-pass Butterworth filter with cut-off frequency of 5 Hz) upper envelope of the EMG signal was used for the analysis. For each muscle and each running condition, the middle 10 stance phases were ensembled averaged and used for analysis.

Agreement between EMG envelopes and the musculoskeletal-modeling-predicted muscle forces was quantified for each participant using the root-mean-squared error (RMSE) and zero-lag cross-correlation. Given the electromechanical delay [7], this approach will focus on the similarity of the waveforms instead of timing of the peaks. Prior to analysis, both EMG and the musculoskeletal modeling-predicted muscle force waveforms were normalized to a 0-1 scale. RMSE was calculated to quantify waveform disagreement with lower values indicating greater agreement. Zero-lag cross-correlation quantified the similarity between EMG and muscle force waveform with temporal shifting.

Differences in agreement metrics among Preferred, Groucho, and Ex Groucho running technique were evaluated using linear mixed models. Running technique was included as a fixed effect and participant was included as a random intercept to account for repeated measures within participants. Type III analysis of variance with Satterthwaite's approximation was used to assess the significance of technique effects. When appropriate, pairwise comparisons between techniques were performed using Bonferroni's correction. Statistical significance was set to 0.05.

### Results

**Figures C1 and C2** depict the ensembled average waveform of EMG and model-predicted force across all the participants and for the seven muscles of interest. The mean and standard deviation of the variables of interest across different muscles and running techniques are presented in **Table C1**. Running condition did not affect agreement between EMG and model-predicted muscle force waveforms. No significant effect of running condition was observed for RMSE ($F(2,19.2) = 0.043$, $p = 0.958$) or for the zero-lag cross-correlation coefficient ($F(2,29) = 0.94$, $p = 0.43$).

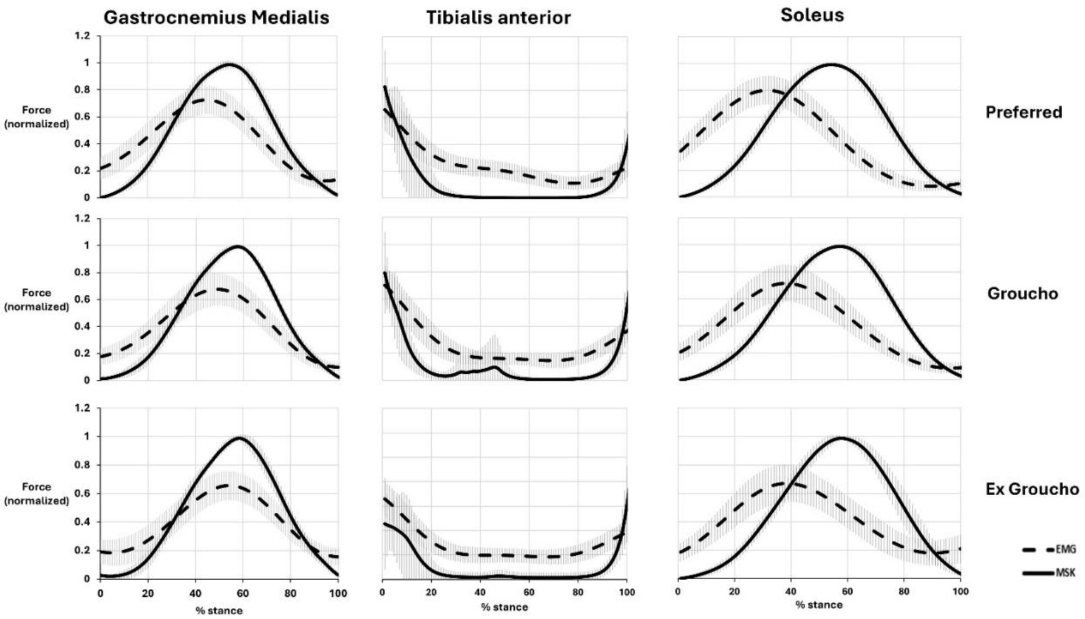


**Figure C 1.** Ensemble-averaged EMG waveforms (dashed lines) and model-predicted muscle forces (solid lines) across all participants during the stance phase. Shaded regions represent ±1 standard deviation. Each column corresponds to a different muscle, indicated by the muscle name at the top of the figure. Each row corresponds to a running technique, indicated by the technique name on the right side of the figure.

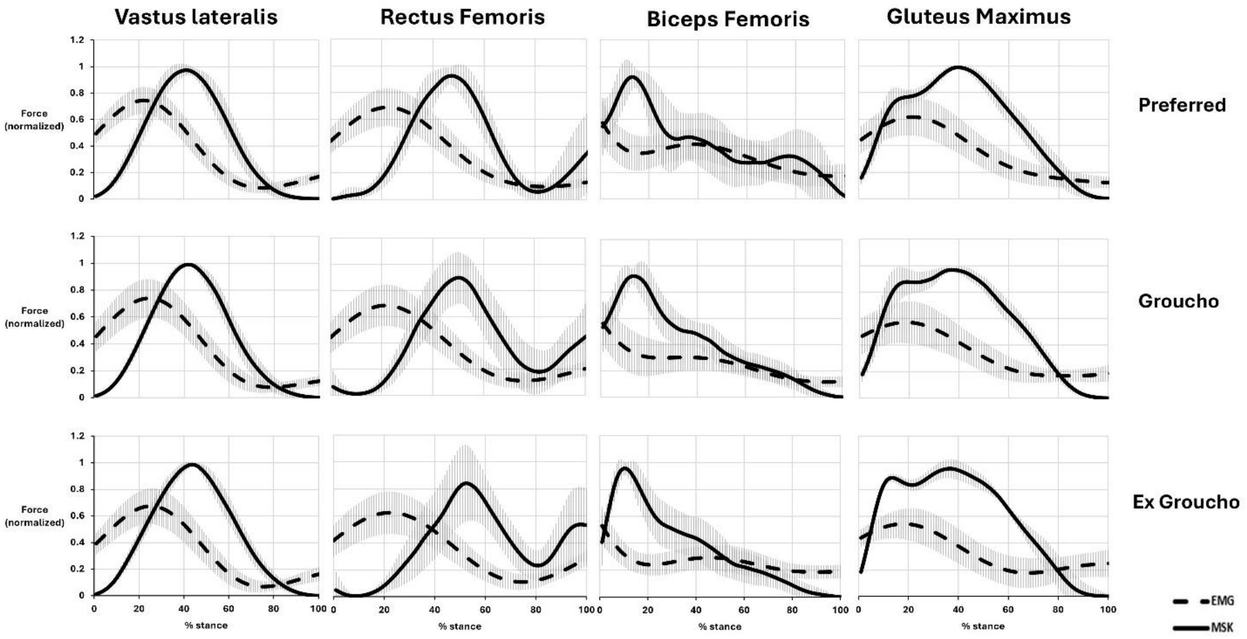


**Figure C 2.** Ensemble-averaged EMG waveforms (dashed lines) and model-predicted muscle forces (solid lines) across all participants during the stance phase. Shaded regions represent ±1 standard deviation. Each column corresponds to a different muscle, indicated by the muscle name at the top of the figure. Each row corresponds to a running technique, indicated by the technique name on the right side of the figure.

**Table C1.** Mean ± standard deviation of the variables of interest for each muscle across different running techniques.

| Muscle | Technique | RMSE | Zero-lag cross-correlation |
|---|---|---|---|
| Gastrocnemius Medialis | Preferred | 0.30± 0.160 | 0.65 ± 0.30 |
| | Groucho | 0.26 ±0.171 | 0.82 ± 0.197 |
| | Ex Groucho | 0.32 ± 0.123 | 0.62 ± 0.263 |
| Tibialis Anterior | Preferred | 0.34 ±0.154 | 0.50 ± 0.238 |
| | Groucho | 0.29 ± 0.108 | 0.58 ± 0.217 |
| | Ex Groucho | 0.16 ± 0.158 | 0.47 ± 0.228 |
| Soleus | Preferred | 0.38 ± 0.084 | 0.37 ± 0.245 |
| | Groucho | 0.36 ± 0.068 | 0.41 ± 0.231 |
| | Ex Groucho | 0.37 ± 0.102 | 0.47 ± 0.272 |
| Vastus Lateralis | Preferred | 0.46 ± 0.76 | 0.31 ± 0.240 |
| | Groucho | 0.46 ± 0.076 | 0.29 ± 0.184 |
| | Ex Groucho | 0.46 ± 0.062 | 0.21 ± 0.151 |
| Rectus Femoris | Preferred | 0.48 ± 0.105 | 0.39 ± 0.241 |
| | Groucho | 0.52 ± 0.040 | 0.22 ± 0.176 |
| | Ex Groucho | 0.50 ± 0.061 | 0.24 ± 0.174 |
| Biceps Femoris | Preferred | 0.40 ± 0.180 | 0.61 ± 0.267 |
| | Groucho | 0.33 ± 0.125 | 0.52 ± .267 |
| | Ex Groucho | 0.35 ± 0.117 | 0.47 ± 0.249 |
| Gluteus Maximus | Preferred | 0.34 ± 0.101 | 0.51 ± 0.199 |
| | Groucho | 0.36 ± 0.088 | 0.43 ± 0.267 |
| | Ex Groucho | 0.40 ± 0.098 | 0.44 ± 0.301 |

### Conclusion

Our findings indicate that running technique did not meaningfully influence the agreement between the EMG-derived muscle activation and musculoskeletal modeling-predicted muscle force waveforms in a manner that would confound our results.
